# High-resolution proteomics identifies potential new markers of Zika and dengue infections

**DOI:** 10.1101/788174

**Authors:** Kristina Allgoewer, Alice Zhao, Shuvadeep Maity, Lauren Lashua, Moti Ramgopal, Beni N. Balkaran, Liyun Liu, Maria T. Arévalo, Ted M. Ross, Hyungwon Choi, Elodie Ghedin, Christine Vogel

## Abstract

Distinguishing between Zika and dengue virus infections is critical for treatment and anticipation of complications. However, existing biomarkers have high error rates. To identify new potential diagnostic signatures, we used next-generation proteomics to profile 122 serum samples from 62 Zika or dengue patients. We quantified >500 proteins and identified 26 proteins that were significantly differentially expressed. These proteins typically function in infection and wound healing, with several also linked to pregnancy and brain. Integrating machine learning approaches, we used 7 proteins to predict ZIKV infection correctly in 72% of the cases, outperforming other tools. The three most predictive proteins were Platelet Factor 4 Variant 1, Fibrinogen Alpha, and Gelsolin. Finally, we showed that temporal changes in protein signatures from the same patient can disambiguate some diagnoses and serve as indicators for past infections. Taken together, we demonstrate that serum proteomics can be highly valuable to diagnose even challenging samples.

## Introduction

Zika virus (ZIKV) and dengue virus (DENV) are closely related Flaviviruses transmitted by the same mosquito vector, *Aedis aegypti*, and with overlapping geographical distributions ^1,2^. While most ZIKV and DENV infections are asymptomatic, they cause a similar immune response and symptoms in the host, including fever and body pain ^1,2^. IgM antibodies typically develop during the first week of illness; however, little is known about the longevity of IgM antibodies following infection ^3^. Neutralizing antibodies, such as IgG, develop shortly after IgM antibodies arise and persist for many years after an infection. Dengue virus-reactive T cells are thought to potentially mediate cross-protection against subsequent ZIKV infections ^1^. In contrast to DENV, ZIKV infections in pregnant women pose a significant risk to the developing embryo, with microcephaly and other adverse outcomes ^44,56^. Therefore, there is a continued need for correct diagnosis of DENV and ZIKV infections as well as determination of past infections. In some geographic areas, diagnosis is purely based on symptoms and endemicity of the virus, which leads to problems due to the shared febrile syndrome. Other affected regions use molecular tests, such as nucleic acid amplification tests (NAATs) of the viral RNA. NAATs are recommended about 7 days after onset of symptoms ^3^. While NAATs are highly sensitive and specific, they produce substantial false-negative and false-positive results. In addition, RNA extraction from the sample can also be difficult due to its instability.

Therefore, NAATs are complemented by antibody-based tests that include Enzyme-Linked ImmunoSorbent Assay (ELISA), and Plaque or Focus Reduction Neutralization Tests. The neutralization tests evaluate the ability of virus specific anti-IgG antibodies to reduce viral infection in cell culture. Dengue is typically diagnosed through IgG and IgM tests in conjunction with geographic location and patient symptoms. The IgM antibody testing should be performed on serum collected less than 7 days after onset of symptoms ^3^. Absence of positive DENV testing in the presence of other symptoms, including pain behind the eyes, often leads to ZIKV diagnosis.

While antibody-based testing is an important diagnostic tool, interpretation of the results is complicated by cross-reactivity of the antibodies leading to false-positives ^37^. In addition, previous infections can impact the assumed time point of the current infection due to antibody longevity. For example, in persons previously infected with, or vaccinated against, a Flavivirus, subsequent infection with another Flavivirus can result in both a diminished IgM response and a rapid increase in neutralizing antibodies against multiple Flaviviruses, which might preclude conclusive determination of which virus was responsible for the person’s most recent infection ^3^. The timing and presence of virus-specific anti-IgM and IgG antibodies is therefore insufficient for diagnosis, in particular in areas where both viruses are endemic. These challenges are particularly critical in pregnant women suspected of ZIKV infections.

For these reasons, highly specific diagnosis of one virus infection over the other is challenging and there is a continued need for additional biomarkers. However, samples are often collected from diverse sites without consistent protocols or note of associated patient data, and sample cohorts can be very small. Further, blood samples can rapidly change in composition depending on storage and processing times. Finally, proteins in serum have enormous abundance differences, e.g. the concentration of serum albumin is an order of magnitude larger than that of other proteins. Even most advanced proteomics studies struggle to identify more than a few hundred proteins ^8,9^. The problem is exacerbated in studies that struggle with obtaining sufficient protein sample or are collected under suboptimal conditions.

We present a study that addresses these challenges by using state-of-the-art proteomics that is particularly amenable to protein mixtures with extreme dynamic ranges ^10^ and analyze a cohort of dengue and Zika patients from Trinidad. We subject the proteomics data to conduct rigorous statistical modeling to remove confounding effects as much as possible and identify differentially expressed proteins with substantial predictive power. Most of the differentially expressed proteins have links to pregnancy and brain. We also gain first insights into proteomic signatures of patients with ambiguous diagnosis which may, in future, contribute to identifying past and co-infections.

## Results

### A cohort of dengue and Zika patients provided complex meta-data

Using high-resolution mass spectrometry with advanced data acquisition techniques, we screened 122 serum samples collected in 2016 and 2017 from a cohort of 62 dengue and Zika patients (**Figure 1A, Supplementary Table S1**). The patients had been recruited from emergency departments in Trinidad. Samples were taken at two different time points after onset of symptoms, i.e. 3-7 and 7-14 days for time points 1 and 2, respectively. For two of the patients (80 and 81), only one time point had been collected.

**Fig. 1.**
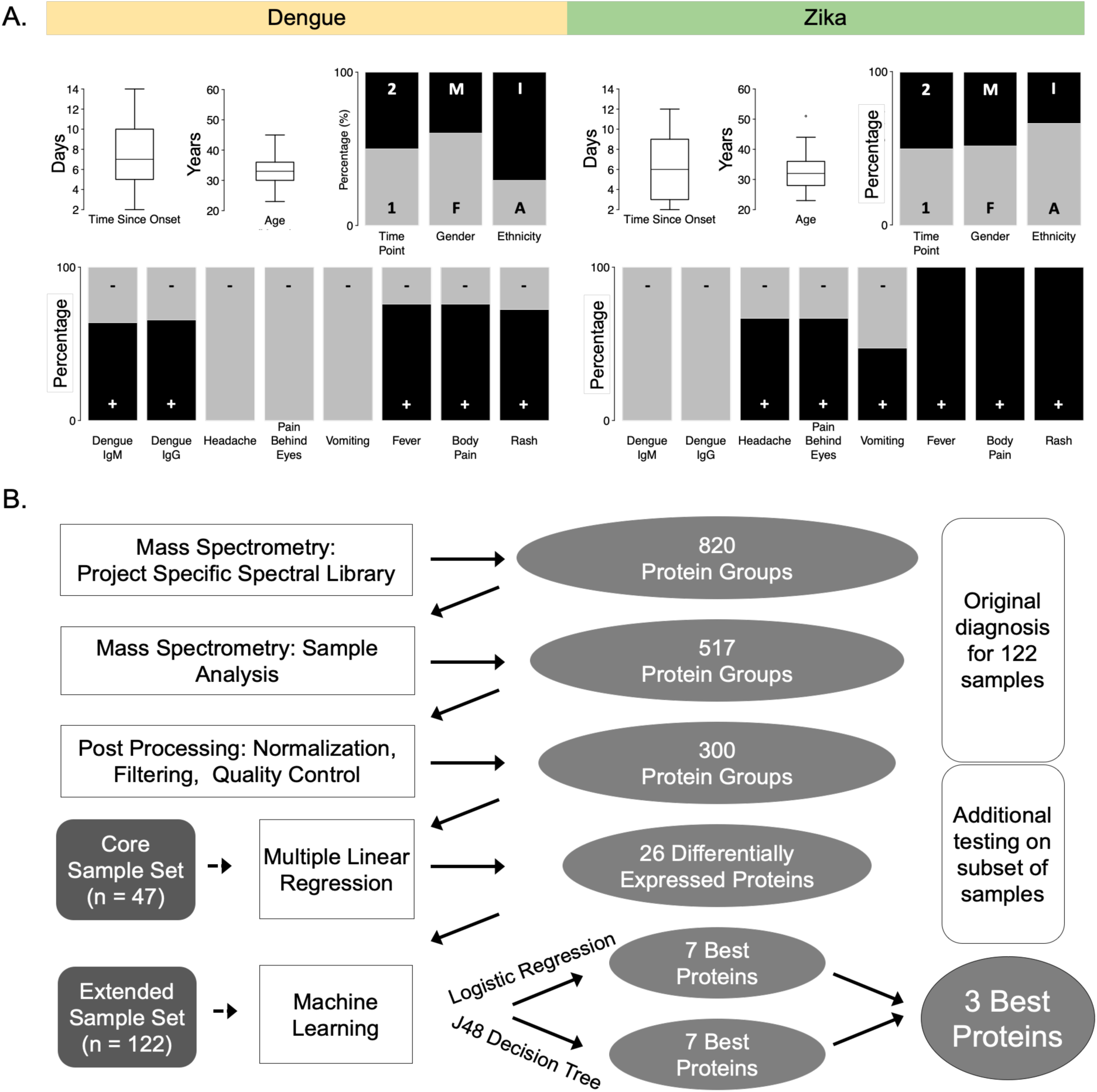
Dengue and Zika patients are from comparable backgrounds. **A.** We collected 122 serum samples from 35 and 27 dengue and Zika patients, respectively. The samples were taken at two time points after onset of symptoms: 3 to 7 days for the first time point (1), 7 to 14 days for the second time point (2). Female (F) and male (M) patients were described as either Afro-Trinidadian (A) or Indo-Trinidadian (I), aged between 23 and 51 years. Diagnosis of dengue and Zika by local doctors was based on a positive (+) or negative (-) dengue IgM/IgG ELISA result and the presence or absence of headache, pain behind the eyes, vomiting, fever, body pain, and rash. **B.** The proteomics and statistical workflow includes: generation of a project specific spectral library of 820 protein groups, detecting 517 protein groups, and retaining 300 protein groups after normalization, filtering, and quality control steps. We then used a core set of 94 samples with complete meta-data to eliminate confounding effects via multiple linear regression, and identify 26 differentially expressed proteins (p-value ≤ 0.01). Employing two different Machine Learning algorithms (logistic regression and J48 decision tree), we predicted the differential diagnosis of dengue and Zika in the extended set of 122 samples, using 26, 7 or 3 of the 26 significant proteins. The three proteins were shared between the best predictors selected from both algorithms.

Patients from the two infection types were similarly distributed with respect to gender, ethnicity (Afro-Trinidadian or Indo-Trinidadian) and age (23-51 years)(**Fig. 1A**). The symptoms (headache, pain behind the eyes, vomiting, fever, body pain, rash) showed biases between the patient groups, but not exclusive classification. Three of the symptoms were only diagnosed in Zika patients and absent in dengue patients: headache, pain behind the eyes, and vomiting.

**Figure 1A** also shows that current diagnostic markers have several inconsistencies. Typically, if patients tested negative in the DENV IgM/IgG ELISA, they were diagnosed as having Zika (**Supplementary Table S1**). However, 9 patients (84, 86, 87, 88, 89, 90, 91, 97, 98) were diagnosed as having DENV, even though positive DENV IgG/IgM were not reported. Patient samples 79-101 had missing IgM/IgG results, but were collected during a dengue outbreak. Due to this heterogeneity in the available meta-data, we performed the modeling on the set of 94 samples (47 patients) with complete meta-data, but tested the predictions on the full set of 122 samples.

### Quantitative proteomics revealed diverse expression patterns across patients

To obtain a quantitative proteomic picture of the patient response to DENV and ZIKV infection, we use our integrated workflow to map 517 groups of indistinguishable protein isoforms across the 122 samples. The lead protein for each protein group provided the name for the group. The heatmap in **Figure 2** presents these data for the 300 protein groups with complete proteomic information (also see **Supplementary Table S2**). Protein abundances ranged over five orders of magnitude, measured as intensity in the mass spectrum. The protein expression patterns were diverse across patients, but show some biases towards diagnosis (dengue vs. Zika patients), gender, and time point of sample collection.

**Fig. 2.**
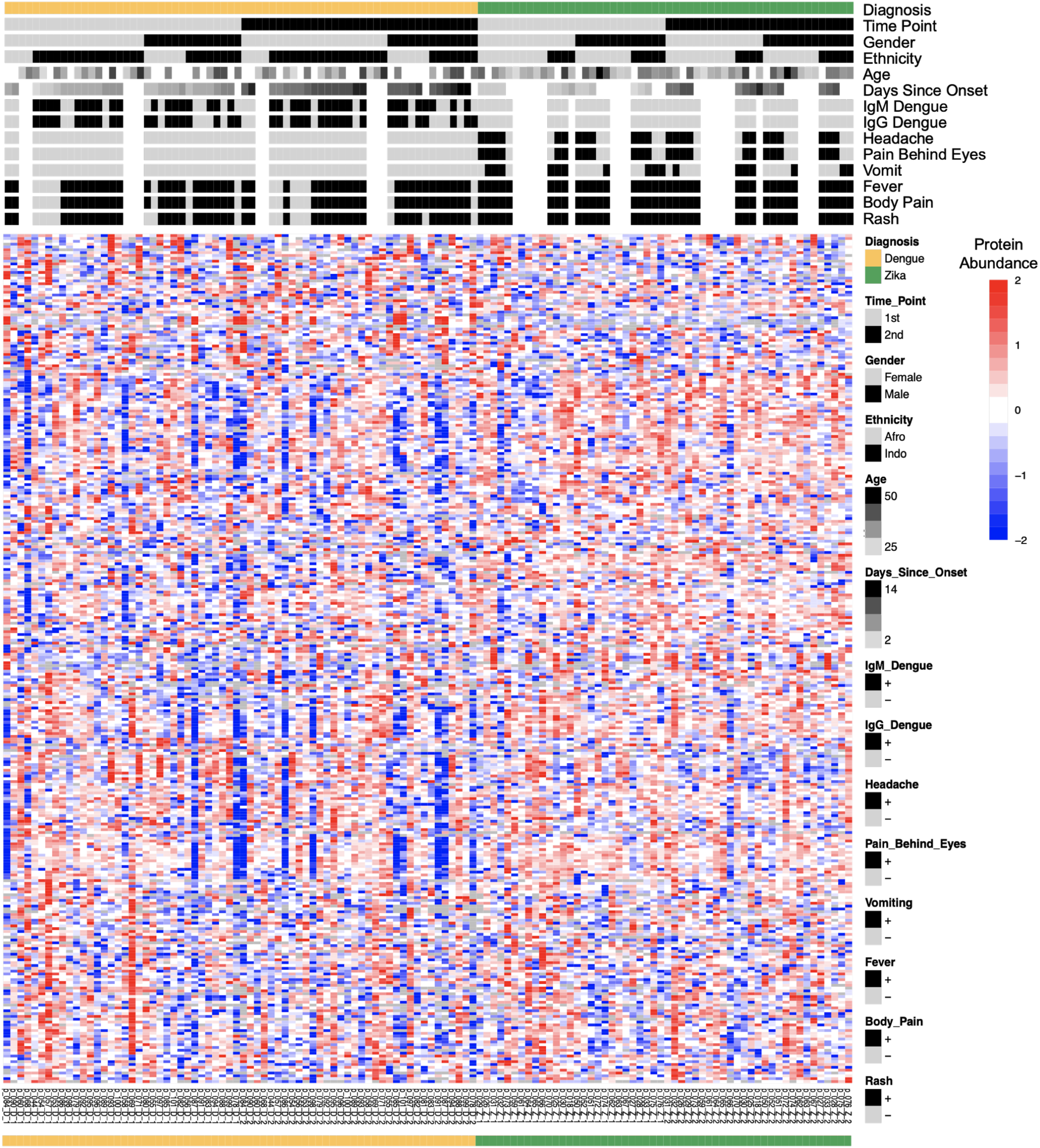
Quantitative proteomics reveals diverse expression patterns across patients. The heatmap shows the normalized and log base 2 transformed protein abundances derived from mass spectrometry. The corresponding meta-data is shown above: diagnosis, sample time point 1 and 2, gender, ethnicity (Afro- and Indo-Trinidadian), age (years), number of days since onset of symptoms, positive (+) or negative (-) dengue IgM and IgG ELISA results, presence (+) or absence (-) of headache, pain behind the eyes, vomiting, fever, body pain, and rash. Missing values in protein data are shown in white.

We examined these biases in patient characteristics in more detail through analysis of the principal components that mark variation in the protein expression matrix. The first five principal components explained 43% of the variation in total (**Table 1, Supplementary Table S3**). The first principal component explained 21% of the expression variation and its patient score correlated significantly with diagnosis (DENV/ZIKV, p-value = 0.002), indicating that most of the expression phenotype is indeed driven by the patient’s pathology. The component also correlated strongly with one clinical variable, ‘vomit’, suggesting it to be an indicator of ZIKV infection (p-value = 0.040). The other components correlated with both diagnosis, detection of the DENV IgM and IgG biomarkers, and some clinical features, such as headache, pain behind the eyes and vomit. These relationships confirmed that a biological signal could be detected in the data but also the presence of several confounding factors.

**Tab. 1.** Variation in protein expression is driven by diagnosis and other clinical variables. The first five principal components from a protein level Principal Component Analysis were correlated with patient meta-data in a linear regression model. The table shows resulting p-values with significant ones (p ≤ 0.05) highlighted in red. The dengue/Zika diagnosis correlates with principal components (PCs) 1 and 2, positive/negative dengue IgM and IgG results with component 2, the presence/absence of symptoms headache and pain behind the eyes with components 2 and 5, and the presence/absence of vomiting with components 1 and 5. The three symptoms--headache, pain behind the eyes, and vomiting--were exclusively diagnosed in Zika patients. Further symptoms, as well as gender, age, ethnicity, and number of days since onset of symptoms, did not correlate with any of the first five principal components. See **Supplement** for correlation of principal components at fragment level.

| Principal component<br>(protein expression variation explained) | Diagnosis | Dengue IgM | Dengue IgG | Head-ache | Pain Behind Eyes | Vomit | Fever | Body Pain | Rash | Gender | Age | Ethnicity | Days Since Onset |
| --- | --- | --- | --- | --- | --- | --- | --- | --- | --- | --- | --- | --- | --- |
| PC1 (21%) | 0.00 | 0.23 | 0.77 | 0.10 | 0.10 | 0.04 | 0.95 | 0.95 | 0.87 | 0.17 | 0.88 | 0.45 | 0.33 |
| PC2 (8%) | $\leq 0.0001$ | 0.00 | 0.01 | $\leq 0.0001$ | $\leq 0.0001$ | 0.08 | 0.63 | 0.63 | 0.90 | 0.76 | 0.84 | 0.61 | 0.96 |
| PC3 (6%) | 0.47 | 0.18 | 0.16 | 0.66 | 0.66 | 0.81 | 0.52 | 0.52 | 0.40 | 0.06 | 0.05 | 0.71 | 0.63 |
| PC4 (4%) | 0.87 | 0.27 | 0.56 | 0.12 | 0.12 | 0.09 | 0.76 | 0.76 | 0.86 | 0.89 | 0.76 | 0.60 | 0.48 |
| PC5 (3%) | 0.14 | 0.30 | 0.25 | 0.03 | 0.03 | 0.02 | 0.24 | 0.24 | 0.25 | 0.96 | 0.71 | 0.40 | 0.55 |

### Isolation of confounding effects identifies 26 differentially expressed proteins

Next, we extracted differentially expressed proteins while accounting for confounding effects. To do so, we focussed on the set of 94 samples with complete meta-data. We removed confounding effects, e.g. age, ethnicity, gender, via multiple linear regression (**Supplementary Table S4**) and identified 26 proteins that were significantly differentially expressed between dengue and Zika patients at a p-value of <0.01 (**Figure 3A**). Most of the proteins (19) were expressed at higher levels in Zika than in dengue. **Supplementary Figure S2** shows differentially expressed proteins at p-value<0.05, confirming the overall trends.

**Fig. 3.**
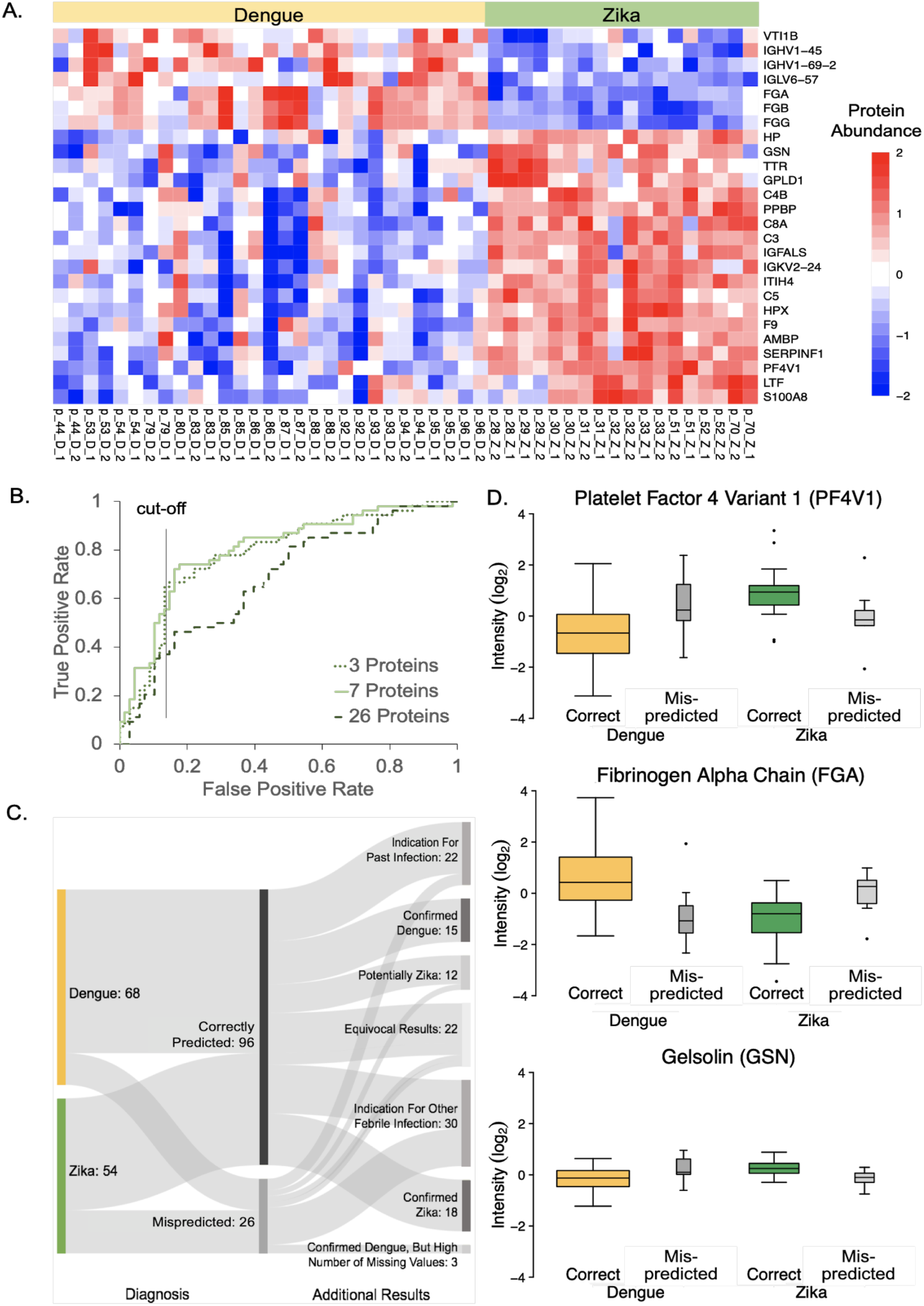
Statistical modeling identifies new potential biomarkers. **A.** Multiple linear regression identifies 26 proteins that are significantly differentially expressed between dengue and Zika patients (p-value ≤ 0.01). The heatmap shows the row-wise clustered and scaled residual abundance data after removing confounding effects. See **Supplementary Figure S2** for the heatmap of 71 differentially expressed proteins at p-value ≤ 0.05. **B.** Using the 26 differentially expressed proteins and two Machine Learning algorithms (logistic regression and J48 decision tree), we predicted the diagnosis for the extended set of 122 samples. Reducing the number to 7 or 3 of the 26 significant proteins improved the true positive rate at the same false positive rate. For Zika samples, we obtained 72% true positives at a 16% false positive rate. **C.** Our predictions are confirmed by outside evidence. Additional experimental testing on a subset of samples (**Supplementary Table S6**) revealed cases of ambiguous diagnosis: dengue samples with an indication for past infections, and some Zika samples with an indication for other febrile infections. The graph evaluates our predictions with respect to the original diagnosis and these additional findings. **D.** Correctly predicted dengue and Zika samples differ in the intensity (log2 transformed) of the 3 best proteins (see **Supplementary Figure S3** for proteins 4 to 7). Mispredicted samples have intermediate intensities.

**Table 2** describes these proteins in their biological roles and known relationships to DENV and ZIKV infections, as well as pregnancy and brain function. **Supplementary Table S5** provides extended descriptions. As expected in serum samples from virus-infected patients, most proteins identified were involved in the immune response and wound repair, as is common in febrile syndromes. The 7 proteins with higher expression values in DENV vs. ZIKV include VTI1B, a protein involved in SNARE mediated vesicle transport. Many of the proteins have known relationships to either DENV and ZIKV infections: interestingly, 5 of the differentially expressed proteins have also been identified in an independent proteomic study comparing dengue patients to healthy controls (C3, C4, HP, HPX, and ITIH4)^11^. Further, with just a few exceptions, all 26 proteins had some relationship to pregnancy, often related to pre-eclampsia, and regulation of brain function (**Table 2, Supplementary Table S5**).

**Tab. 2.** Differentially expressed proteins may present new biomarkers of disease. The table shows the 26 differentially expressed proteins (p-value ≤ 0.01) and their known connections to dengue and Zika as well as to brain and pregnancy related research. These proteins were identified by removing confounding effects through multiple linear regression in a set of 94 samples with complete meta-data.

| Protein | Gene | Function | Levels in Zika vs dengue serum | Relationship to |  |  |  |
| --- | --- | --- | --- | --- | --- | --- | --- |
|  |  |  |  | Dengue | Zika | Pregnancy | Brain |
| **Platelet factor 4 variant 1** | PF4V1*,** | Inhibitor of angiogenesis and of endothelial cell chemotaxis in vitro | Up | yes | yes | yes | yes |
| **Fibrinogens** | FGA*,**FGB, FGG | Primary components of blood clots in early stages of wound repair | Down | yes | yes | yes | yes |
| **Gelsolin** | GSN*,** | Binds to the plus ends of actin monomers or filaments, preventing monomer exchange | Up | n/a | n/a | yes | yes |
| **Inter-alpha-trypsin inhibitor heavy chain H4** | ITIH4** | Inflammatory responses to trauma | Up | yes | yes | yes | yes |
| **Complement Components** | C3**, C4B, C5, C8A | Protection of the host from infection/inflammation by recruiting and enhancing phagocytosis | Up | yes | yes | yes | yes |
| **Transthyretin** | TTR** | Probably transports thyroxine from the bloodstream to the brain | Up | yes | n/a | yes | yes |
| **Pigment epithelium-derived factor** | SERPINF1** (PEDF) | Neuronal differentiation in retinoblastoma cells, inhibitor of angiogenesis | Up | n/a | n/a | yes | yes |
| **Lactotransferrin** | LTF | Stimulates VEGFA-mediated endothelial cell migration and proliferation | Up | yes | yes | yes | yes |
| **Platelet basic protein** | PPBP | Chemoattractant and activator of neutrophils, stimulates various cellular processes including DNA synthesis, mitosis, glycolysis, intracellular cAMP accumulation, prostaglandin E2 secretion, and synthesis of hyaluronic acid and sulfated glycosaminoglycan | Up | yes | yes | yes | yes |
| **Immunoglobulins** | IGHV1-45, IGHV1-69-2, IGKV2-24, IGLV6-57 | Antigen recognition | Down | yes | yes | yes | yes |
| **Protein AMBP, alpha-1-microglobulin/ bikunin precursor** | AMBP | Processed alpha-1-microglobulin may be involved in the regulation of inflammatory processes | Up | yes | n/a | yes | yes |
| **Haptoglobin** | HP | Captures and combines with free plasma hemoglobin to allow hepatic recycling of heme iron | Up | yes | n/a | yes | yes |
| **Hemopexin** | HPX | Bins heme and transports it to the liver for breakdown and iron recovery | Up | yes | n/a | yes | yes |
| **Coagulation factor IX** | F9 | Blood coagulation | Up | yes | n/a | yes | n/a |
| **Insulin-like growth factor-binding protein complex acid labile subunit** | IGFALS | Protein-protein interactions that result in protein complexes, receptor-ligand binding or cell adhesion | Up | n/a | n/a | yes | yes |
| **Phosphatidylinositol-glycan-specific phospholipase D** | GPLD1 | Hydrolyzes the inositol phosphate linkage in proteins anchored by phosphatidylinositol glycans | Up | n/a | n/a | yes | yes |
| **Protein S100-A8** | S100A8 | Inflammatory processes and immune response, induces neutrophil chemotaxis and adhesion | Up | n/a | n/a | yes | yes |
| **Vesicle transport through interaction with t-SNAREs homolog 1B** | VTI1B | Vesicle transport pathways through interactions with t-SNAREs | Down | n/a | n/a | n/a | yes |

### Machine learning highlights potential new diagnostic markers

To select a sparse panel of candidate markers and evaluate their diagnostic performance, we extracted the proteomic profiles for the 26 differentially expressed proteins for all 122 samples of the extended dataset and used two different algorithms to predict the diagnosis. We evaluated the true and false positive rates of the two algorithms using the primary diagnosis made when the samples had been acquired (**Figure 3B**). For each algorithm, we extracted a set of 7 proteins with the highest predictive capabilities: 3 proteins were shared between the outputs of the two algorithms, which we defined as our ‘Best Proteins’ (**Figure 1**).

Predictive performance was better with the core proteins than with the 26 differentially expressed proteins. While the 26 proteins predicted with a true-positive rate of 0.38 across the 122 samples, the 7 and 3 core proteins predicted with a true-positive rate of 0.72 and 0.71, respectively (false positive rate = 0.16; **Figure 3B**). We examined the mispredictions in more detail, using data from additional experimental testing of samples, such as ZIKV IgM and IgG that confirmed or modified the original diagnosis (**Figure 3C, Supplementary Table S6**). All of the 18 ZIKV samples for which diagnosis was confirmed by additional testing had been correctly classified using our approach. For DENV, 15 out of 18 confirmed samples had been correctly predicted, while the 3 mispredicted samples had many missing values.

Next, we examined the most predictive proteins in more detail to verify their putative roles during the response to infection. In particular, we extracted the 3 ‘Best proteins’ identified by both algorithms as having the strongest power with respect to predicting diagnosis: Fibrinogen Alpha Chain (FGA), Gelsolin (GSN), and Platelet Factor 4 Variant 1 (PF4V1). The distributions of theiir protein expression levels are shown in **Figure 3D. Supplementary Figure S3** shows additional examples of the best potential biomarkers. FGA and other fibrinogens are components of blood clots and involved in early wound repair. FGA is one of the few proteins with lower expression levels in Zika compared to dengue, although plasma fibrinogen concentrations are known to be repressed in dengue patients compared to control ^12^. Furthermore, DENV and its antibodies can directly influence the fibrinolytic pathway ^13^. In comparison, a Zika patient with severe liver injury and coagulation disorders showed altered blood levels of fibrinogen and fibrinogen degradation products ^14^. FGA is also linked to embryonic development and remyelination of the nervous system ^15,16,17^.

PF4V1 is an inhibitor of angiogenesis and expressed at higher levels in Zika than in dengue patients. Consistently, it has been found down-regulated in platelets from dengue patients compared to control ^18^, as high levels can promote rapid replication and propagation of the virus ^19^. Indeed, the platelet count is one of the diagnostic indices used to distinguish between DENV and ZIKV infections ^20^. GSN (Gelsolin) binds to the plus ends of actin monomers or filaments. Similar to FGA and PF4V1, it is also linked to pregnancy and brain function: its maternal plasma levels are upregulated through pregnancy-related hormones and in the brain during aging ^21,22^.

### Time-resolved proteomics illustrates patterns for patients with potential past infections

Finally, we examined the samples with ambiguous diagnosis for signatures of past infections, i.e. with Flavi- or other viruses. To do so, we applied our computational pipeline to the core set of samples, but labeled the samples as arising from either unambiguous or ambiguous diagnosis. We defined unambiguous diagnosis as those cases that were confirmed by additional serology. We defined ambiguous diagnosis for two patient groups. First, we used patient samples with an original dengue diagnosis, but positive additional testing for Zika IgG or IgM in at least one time point. In total, 17 patients (34 samples) qualified. Second, we used patient samples with an original Zika diagnosis but with negative Zika serology as Zika patients; this applied to 15 patients (30 samples). Note that each group of ambiguous cases likely contains several false-positives arising from cross-reactivities of the respective antibodies.

We then applied our computational pipeline to remove confounding factors and identified 6 proteins with significant differential expression between the samples with unambiguous and ambiguous diagnosis (pvalue<0.01; THBS1, AHSG, DHX9, IGLC2, TFRC, VTI1B; **Supplementary Figure S4**). Three of the factors were expressed at higher levels in ambiguous than in unambiguous cases - AHSG, TFRC, DHX9 - and are involved in inflammation, innate immunity, and immunodeficiency which are components of past infections ^23,24,25,26^.

Further, we aimed at identifying temporal signatures of past infections. To do so, we examined the cases of ambiguous and unambiguous diagnosis for expression changes across the two measurement time points. We first compiled average protein expression profiles of the correctly predicted dengue and Zika patient samples which had been confirmed by additional serological testing (**Figure 4A, Supplementary Table S6**). These average profiles showed only small changes in protein expression between the two time points, but distinct differences in expression levels between dengue and Zika. Each infection had a unique signature. We then calculated the average protein expression profiles for patients with ambiguous diagnosis (**Figure 4A**). The average expression levels were very similar across proteins or between time points, without distinct differences.

**Fig. 4.**
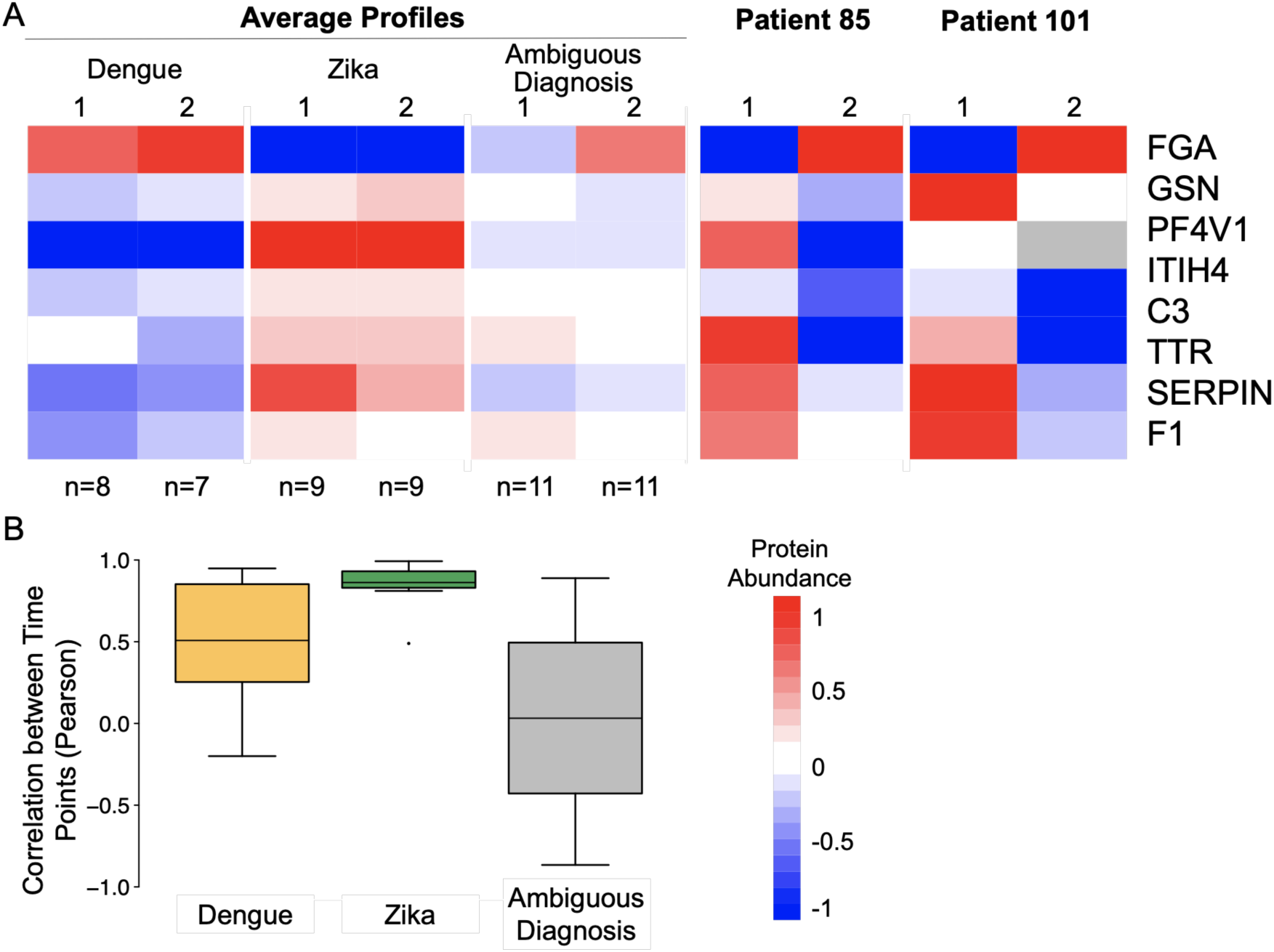
Potential cases of past infections show distinct expression patterns between consecutive time points. **A.** The average dengue and Zika profiles show strong expression differences amongst the 7 core proteins and high similarity between the two measurement time points. These expression differences are not observed for the average profiles of samples with ambiguous diagnosis and evidence for dual infections. Note that dual infections have not been confirmed. When plotting individual patient samples with ambiguous diagnosis (Patient 85 and patient 101), expression differences become apparent: within approximately one week between the first and second measurement point both patients switch from a Zika-like to a dengue-like expression pattern. **B.** This trend is confirmed when examining average correlation of protein expression profiles from the two time points across groups of patients: patients with correctly predicted, unambiguous, single infections show a much higher temporal correlation in protein expression than patients with an indication for past infections.

However, when we examined individual patients with ambiguous diagnoses, such as patients 85 and 101, we observed a striking trend (**Figure 4A**). Neither of the two patients showed ‘vomit’, ‘pain behind the eyes’ or ‘headache’ as a symptom, which are exclusive characteristics for acute ZIKV infection, and they had both been originally diagnosed with dengue at the time of sample collection. Additional serological testing could not confirm the diagnosis. Indeed, both patients showed protein expression patterns similar to that of Zika in the first time point, contrasting the original diagnosis. Only in the samples from the second time point - collected about 7 days later -- the protein expression patterns were similar to that of unambiguously diagnosed dengue patients. The change in expression was particularly striking for FGA, ITIH4, and C3. Both patients may have had previous ZIKV infections, which affected the response to the acute DENV infection with respect to temporal switch in protein expression profiles. This interpretation is consistent with DENV and ZIKV outbreaks in the region at the time of, and prior to, sample collection (**Methods**).

Finally, we confirmed the temporal disconnect between expression patterns for the entire set of patients with ambiguous diagnosis (**Figure 4B**). To do so, we examined the correlation between the protein expression signatures of the consecutive measurement time points for each patient. The correlation of expression values was low for patients with evidence for past infections and much higher for unambiguously diagnosed dengue or Zika patients (**Figure 4B**). Therefore, the response to an acute infection (onset of symptoms) may at first be dominated by the signature characteristic for past infections, but then switch to countering the current virus.

## Discussion

Our study provides a comprehensive proteomic analysis of a cohort of 62 dengue and Zika patients. Our integrative proteomic and statistical approach enabled us to overcome the substantial challenges arising from heterogeneity in meta-data and sample collection, and to identify new candidates for protein biomarkers of these infections. We quantified >500 proteins across the serum samples, with the complete dataset consisting of 300 proteins. We extracted 26 differentially expressed proteins from these data. Most of these differentially expressed proteins have links to pregnancy and brain function. We validated these proteins through comparison with the literature: 5 of the 26 proteins had been identified in serum proteomes from DENV infected patients compared to healthy controls in India ^11^: Haptoglobin, Complement C3 and C4, Hemopexin, and Inter-alpha-trypsin inhibitor heavy chain H4.

Using proteomic data from the 26 differentially proteins or sets of 7 most predictive proteins, we were able to predict the dengue and Zika diagnosis with high sensitivity and specificity, achieving 72% true positive Zika identifications at a 16% false positive rate. These results outperformed other predictions based on the presence of viral proteins in blood samples ^8^. We extracted 3 ‘Best proteins’ -- FGA, PFV4F1, and GSN -- which occurred at the intersection of these predictive approaches. Literature confirmed the potential of these proteins as new biomarkers: PFV4F1, for example, has strong links to DENV propagation ^19^. Further, all three proteins showed links to pregnancy and brain function, which could help explain adverse effects of ZIKV infections.

Four of the differentially expressed proteins were linked to the complement system (C3, C4B, C5, C8A), with C3 being one of the most predictive core proteins. The complement system has an ambivalent role in Flavivirus infection: it can be protective by limiting viral replication, or contribute to disease severity when excessively activated and causing an exacerbated inflammatory response ^27^. Its occurrence amongst the differentially expressed proteins might link to the increased activity of the complement systems in DENV infections ^28,29^.

Finally, we investigated the proteomic data for diagnostic signatures of possible past infections with dengue/Zika or other viruses. Some geographical areas have a high presence of both Flaviviruses ^30^, and Trinidad - the origin for the cohort analyzed here - witnessed a DENV outbreak in 2015/2016, simultaneously with reports on increased ZIKV occurrences in 2016/2017 (http://paho.org).

To test for proteomic signatures of past infections, we divided the cohort into unambiguously diagnosed patients and those with ambiguous test results. When applying our computational pipeline to remove confounding factors, we identified 6 proteins with significant differential expression (p-value<0.01, **Supplementary Figure S4**). Amongst these proteins was Fertuin-A (Alpha-2-HS-glycoprotein) which had higher expression levels in cases of ambiguous than unambiguous diagnosis. Fertuin-A has not only been identified as an interactor of DENV protein NS1 ^31^, but also is a known marker of inflammation during acute phase infections ^24,32,33^. Its anti-inflammatory roles ^34^ might explain its higher levels in patients with ambiguous diagnoses as these samples are enriched for patients with prior infections and co-infections that stimulate inflammation.

Further, we identified clear proteomic patterns across the two measurement time points obtained for each patient: patients with unambiguous, single infections showed temporally consistent expression of the core proteins. In contrast, patients with mixed diagnoses show clear temporal evolution in the protein expression profiles over the course of the ∼1 week between the two different time points. While intriguing and based on rigorous statistical filtering, these protein expression signatures of patients with ambiguous diagnoses are based on small cohorts and need to be subjected to further investigation before serving as markers of past infections.

## Methods

### Sample collection

Patient serum samples were collected from Trinidadian dengue and Zika patients in 2016 and 2017. Female and male patients were described as either Afro-Trinidadian or Indo-Trinidadian, aged between 23 and 51 years. Diagnosis of dengue and Zika by local doctors was based on positive or negative DENV IgM/IgG results using a rapid, dual IgM/IgG test (Panbio Dengue Duo Cassette) and the presence or absence of symptoms such as headache, pain behind the eyes, vomiting, fever, body pain, and rash. The three symptoms (headache, pain behind eye, and vomit) were presumably used by local doctors to distinguish Zika from dengue. **Supplementary Table S1** lists meta-data for all samples. **Supplementary Table S6** lists results for additional serology tests with antibodies against ZIKV IgM and IgG in Focus Reduction Neutralization Tests. These additional tests were conducted at a later time point. The Pan American Health Organization reports total incidents for Trinidad/Tobago as: dengue: 2014: 128; 2015: 1,687; 2016: 1,801; 2017: 300; 2018: 123 and Zika: 2014: na; 2015: 0; 2016: 722; 2017: na; 2018: na (https://www.paho.org).

### Sample preparation

We resuspended serum samples (∼75µg) with 0.1% Rapigest (Waters) in 100 mM ammonium bicarbonate (Sigma-Aldrich) and incubated for 5 minutes at 95°C to facilitate protein denaturation. It was then reduced with 5 mM dithiothreitol (Sigma-Aldrich) for 30 mins at 60°C, followed by alkylation with 15 mM iodoacetamide (Sigma-Aldrich) at room temperature for 30 minutes in the dark. We digested the samples overnight using sequencing grade modified porcine trypsin (w/w ratio 1:50) (Sigma-Aldrich) on a thermomixer at 37 °C, 200 RPM (Eppendorf). Rapigest surfactant was cleaved by incubating samples with ∼200 mM HCL (Sigma-Aldrich) for 30 min at 37 °C. We desalted digested protein samples on C18 spin tips (Thermo Fisher Scientific) and dried the peptides under vacuum. The dried peptides were resuspended in 5% acetonitrile, 0.1% formic acid (Sigma-Aldrich). The peptide concentration was measured using a fluorometric peptide quantification kit (Thermo Fisher Scientific).

For the mass spectrometry analysis, we constructed a pooled quality control (QC) sample which contained aliquots of 10 randomly chosen samples. To construct the spectral library, we pooled aliquots from all 124 samples and fractionated the mixture using high-pH reversed-phase high performance liquid chromatography on an Agilent 1200 Infinity Series HPLC with a phenomenex Kinetex 5 u EVO C18 100A column (100 mm × 2.1 mm, 5 mm particle size). Mobile phase A contained 20 mM ammonium formate, and B contained 90% acetonitrile and 10% 20 mM ammonium formate. Both buffers were adjusted to pH 10. Peptides were fractionated using a linear 70 min 0 to 40% acetonitrile gradient at a 100 µl/min flow rate. Eluting peptides were collected into 2 min fractions. We combined fractions to 20 samples for mass spectrometric analysis. The volume of re-combined fractions was reduced using an Eppendorf Concentrator Vacufuge Plus and suspended in HPLC-grade water containing 5% acetonitrile and 0.1% formic acid.

### Mass spectrometry

#### Instrumentation

We used an EASY-nLC 1000 coupled on-line to a Q Exactive High Field mass spectrometer (both Thermo Fisher Scientific) for chromatography and mass spectrometry, respectively. Buffer A (0.1% formic acid in water) and buffer B (80% acetonitrile, 0.1% formic acid) were used as mobile phases for gradient separation. Separation was performed using a 50 cm x 75 µm i.d. PepMap C18 column (Thermo Fisher Scientific) packed with 2 µm, 100 Å particles and heated to 55 °C. We used a 155 min segmented gradient of buffer A to buffer B at a flow rate of 250 nl/min as follows: 2 to 5% buffer B for 5 min, 5 to 25% buffer B for 110 min, 25 to 40% buffer B for 25 min, 49 to 80% buffer B for 5 min and 80 to 95% buffer B for 5 min. Buffer B was held at 95% for another 5 min.

#### Data independent acquisition (DIA)

All serum samples were randomized in their run order and examined in batches of 4-8 samples. The QC sample was run before and after each batch. **Supplementary Table S1** details the run order. All samples were analyzed as follows: a full-scan MS was acquired in the Orbitrap with a resolution of 120,000, scan range of 350–1650 m/z, maximum injection time of 100 ms, and an Automatic Gain Control (AGC) target of 3e6. Subsequently, 17 DIA variable windows were acquired in the Orbitrap with a resolution of 60,000, AGC target of 1e6, and maximum injection time in auto mode.

#### Data dependent acquisition (DDA)

For the spectral library we analyzed the fractionated samples in DDA mode. Full MS scans were acquired with a resolution of 120,000, an AGC target of 3e6, a maximum injection time of 100 ms, and scan range of 375 to 1500 m/z. Following each full MS scan, data-dependent high-resolution HCD MS/MS spectra were acquired with a resolution of 30,000, AGC target of 2e5, maximum injection time of 50 ms, 1.5 m/z isolation window, fixed first mass of 100 m/z, and NCE of 27 with centroid mode.

### Computational analysis to identify differentially expressed proteins

#### Primary analysis

All 144 DIA samples consisting of 124 patient and QC samples were analyzed using Spectronaut (Software Version: 12.0.20491.0.25225) against the project-specific spectral library using default settings. The library was obtained using the Pulsar search engine within Spectronaut with default settings that included Trypsin/P digest, peptide length of 7-52 amino acids, and up to 2 missed cleavages. The FASTA file was downloaded from UniProt on 2/15/2018 and contained 93,798 entries including protein isoforms. Two samples failed in their data acquisition.

#### Secondary analysis

We used in-house R scripts to process Spectronaut output, e.g. with respect to deriving a unique fragment identifier and removing replicate fragment ions. We also removed fragment entries with <50 intensity and any faulty samples (e.g. patient 48 which did not contain protein). We filtered for fragments with <40% missing values across samples. We removed a batch effect caused by replacement of the chromatographic column by fitting lowess curve (f=0.25) through each fragment and subtracting the fitted value from each sample within each batch (**Supplementary Figure S1**). We then set the average to a common overall average peak area value. We then removed all values that remained 3 or more standard deviations away from the median.

#### Tertiary analysis

Principal Component Analysis was conducted with the prcomp() function using log base 2 transformed and mean centered expression data with missing values imputed with the impute.knn function with default settings. **Supplementary Table S3** shows the correlation of the principal components with meta-data at the fragment and protein level.

We then used mapDIA ^5^ to derive protein-level abundance values from fragment ion intensities and added a 2,000 count to all values, resulting in 300 protein groups with high-quality quantitation (**Supplementary Table S2**). To discover differentially expressed proteins adjusting for confounders, we used in-house R-scripts to develop a multiple linear regression model for the core set of samples with complete meta-data, accounting for correlations with Diagnosed_Type, Dengue IgM, Gender, Ethnicity, Days_since_onset, and Age as listed in **Supplementary Table S4**. We excluded the Dengue IgG information due to its high correlation with the Dengue IgM data (identical status). We subtracted the median values for continuous variables (Age, Days_since_onset) to ensure consistent scaling. Multiple linear regression using the lm() function in R resulted in p-values associated with each protein (**Supplementary Table S3**). Clustering was performed with the hclust() function with default settings. Heatmaps were produced with the pheatmap() function.

### Diagnostic marker selection and performance evaluation

We extracted the normalized protein abundance counts for the 26 proteins with significant expression differences between dengue and Zika patients to predict diagnosis for the complete set of 122 patient samples. To do so, we used the WEKA machine learning environment and tested various built-in algorithms for performance, identifying Logistic Regression and J48 decision tree as best. Performance was evaluated by the percentage correctly classified samples during 10-fold cross-validation. After modeling with all 26 proteins, we gradually reduced the number of proteins by using only those with an absolute coefficient of at least 0.5, resulting in identifying 7 proteins with the best results in both best-performing algorithms. The sets intersected with 3 proteins, which we designated as the 3 Best Proteins (**Figure 1**).

### Data availability

All raw mass spectrometry data are deposited in PRIDE including primary Spectronaut output files. R scripts are deposited in github. Exact references will be available upon publication of the manuscript.

## Acknowledgements

K. A. acknowledges funding by the pre-doctoral fellowship by the German Academic Exchange Service (DAAD, “Jahresstipendium für Doktorandinnen und Doktoranden bei bi-national betreuten Promotionen”). H.C. acknowledges funding by the National Medical Research Council of Singapore (NMRC-CG-M009). T.M.R. is also supported, in part, by the Georgia Research Alliance. C.V. acknowledges funding by the National Institutes of Health (R35 GM127089) and the Zegar Family Foundation for Genomics Research. E.G. and C.V. were supported by National Institutes of Health U01 AI111598. We thank Ileana Cristea for useful discussions.

## Author contributions

K.A. conducted all computational analyses and wrote the manuscript. A.Z. and S.M. conducted experiments. L.L., M.R., B.N.B., L.L, M.T.A., and T.M.R. acquired and managed samples. T.M.R., E.G., and C.V. lead the project and wrote the manuscript. H.C. contributed to the analyses and manuscript writing. All authors edited the manuscript.

